# Rigorous Validation of Paw Preference Using Three Complementary Behavioral Assays in Sprague Dawley Rats

**DOI:** 10.64898/2026.02.08.704691

**Authors:** Dipesh Pokharel, Khoi Le, Dilshan Beligala, Thyagarajan Subramanian, Kala Venkiteswaran

## Abstract

Paw preference, or handedness, is a widely studied behavioral trait used to assess lateralization and motor function in rodents. This study aimed to determine the consistency and reliability of three commonly used behavioral tests to rigorously assess paw preference: the Collins Test, the Staircase Test, and the Pawedness Trait Test. Thirty Sprague Dawley rats (12–48 weeks; 20 females, 10 males) were subjected to all three behavioral tests. Paw uses were recorded, and the laterality index was calculated for each test. Additional cohorts of younger rats (6–9 weeks; 45 females, 45 males) and older rats (12–48 weeks; 38 females, 45 males) were tested to assess the effects of age and sex on paw preference. ANOVA, Fleiss’ and pairwise Cohen’s Kappa were used for statistical analysis. All three tests yielded comparable measures of paw preference (ANOVA, p = 0.801). Substantial inter-test agreement was demonstrated by Fleiss’ kappa (κ = 0.761, p = 3.93 × 10□^12^). Paw preference did not significantly vary by age or sex, and the distribution of left, right, and ambidextrous preference categories aligned with existing literature. The Collins, Staircase, and Pawedness Trait Tests provide consistent, reliable assessments of paw preference in Sprague Dawley rats. These validated behavioral assays can serve as essential tools for preclinical research, including but not limited to models of motor asymmetry observed in stroke, cerebral palsy, traumatic brain injury, and language lateralization, as well as neurodegenerative diseases.

**Highlights:**

- Rigorously validated paw preference in Sprague Dawley rats using three commonly used behavioral tests.
- Demonstrated strong inter-test agreement across Collins, Staircase, and PaTRaT (κ = 0.761)
- Showed that paw preference remains stable across age and sex in large cohorts
- Applied standardized LI thresholds to enable cross-test comparability

## 1. Introduction

Lateralization is a fundamental feature of brain organization, evident across both vertebrate and invertebrate species.^1^ Studies in primates, birds, reptiles, canines, rodents, and even insects demonstrate that lateralized behaviors are a conserved feature of nervous system function.^2-9^ In rodents, paw preference provides an analogue to human handedness and serves as a practical behavioral index of hemispheric specialization.^9-11^ This trait holds particular relevance for modeling neurodegenerative diseases such as Parkinson’s disease (PD), Huntington’s disease, and amyotrophic lateral sclerosis (ALS), which frequently manifest with asymmetric motor symptoms.^12-16^ Although recent studies have explored alternate laterality measures such as head turning asymmetry and pasta matrix task, we focused our investigation specifically on reaching-based paw preference under controlled laboratory conditions as a rigorous index of motor lateralization in rats.^17, 18^

Reaching tasks occupy a central role in neurobehavioral research because they enable quantitative assessment of motor lateralization in relation to underlying brain asymmetries.^10, 13, 19-21^ These paradigms have been extensively applied not only in rodents but also in primates, dogs, and birds, underscoring their comparative and translational value.^17, 22-25^ Methodological refinements have emphasized the importance of species-specific anatomy and task design, to ensure reliable assessment of accounting for lateralized motor function.^9, 17^ Rat paw preference has been associated with molecular, functional, and behavioral indices of cerebral asymmetry, reinforcing its value as an accessible preclinical measure.^20, 21, 26-30^.

It is well known that many neurological disorders have an onset in one hemisphere of the brain resulting in symptoms only on one side of the body. For example, stage-1 Parkinson’s disease is defined by hemiparkinsonism. Previous animal studies have suggested that disease vulnerability may be associated with paw preference in rodents.^13, 28, 31^ Clinical studies also suggest that the onset of parkinsonism in the right hemisphere in right-handed (left hemisphere dominant) is associated with worse prognosis for more rapid progression of motor symptoms and cognitive decline. Similar pathological correlates of brain lateralization to disease onset have been noted in Huntington’s disease, ALS, and multiple sclerosis.^12, 32-35^ Therefore, it is critical to document paw preference in rodent models of neurological disorders such PD, Huntington’s disease, ALS, and multiple sclerosis. Differences in task duration, apparatus design, food restriction, and environmental conditions can significantly influence paw preference outcomes, leading to inconsistent findings.^9, 36-39^

To mitigate these methodological inconsistencies, several standardized assays have been developed. The Collins Test, introduced by Collins (1968), involves the forced use of a single forepaw to retrieve pellets through a narrow slit, providing a direct measure of paw dominance.^40^ The Staircase Test Montoya et al., (1991) requires skilled retrieval from bilaterally placed stepped platforms, allowing precise assessment of lateralized forelimb use and fine motor coordination.^41^ In contrast, the Pawedness Trait Test (PaTRaT) Cunha et al., (2017) evaluates spontaneous paw use in a semi-naturalistic setting, minimizing stress and capturing unforced behavior.^42^ This study directly compares these three widely used tests to rigorously measure paw preference in rats. Establishing reliability across these assays will enhance their utility for modeling motor asymmetry and expand their application to studies investigating asymmetric neurodegenerative and neurological disorders.^12, 14, 15, 32, 43^

## 2. Materials and Methods

### 2.1 Animals

Male and female Sprague Dawley rats (6–48 weeks) were housed two per cage in an AAALAC-accredited facility under a 12:12 h light/dark cycle (lights on 07:00, off 19:00). Cages were enriched with nesting material and PVC tubes. Food and water were provided with *ad libitum*, except during mild food restriction prior to behavioral testing. All rats were task-naïve at study onset. Behavioral testing was conducted during the light phase (09:00–15:00), consistent with standard practice, and animals were habituated to the testing room for ≥30 min before each session.

To enhance motivation, rats received 15 grams of chow per rat for 24hrs prior to testing, maintaining body weight at ≥90% of free-feeding levels. Outside of restriction periods, food and water were freely available. All procedures were approved by the University of Toledo IACUC and complied with the Animal Welfare Act and the National Research Council’s *Guide for the Care and Use of Laboratory Animals*. Behavioral testing was performed by the same trained experimenters, who was blinded to animal age and sex during scoring.

Thirty rats (20 females, 10 males; 12–48 weeks) formed the core inter-test cohort and completed all three assays (Collins, Staircase, PaTRaT). A minimum interval of 10 days separated tests, during which animals were returned to their home cages with *ad libitum* access to food and water. Test order was fixed (Collins → Staircase → PaTRaT) to maintain consistency. Data indicated that the 10-day interval was sufficient to prevent fatigue, training, or stress effects.

Additional cohorts were assessed using only the Collins Test to examine broader age- and sex-related effects (young: n=90; old: n=83; males: n=93; females: n=80). The Collins Test was selected due to its reliability, feasibility for high-throughput screening, and widespread historical use.^40, 44-47^ Full cohort details are provided in Supplemental Table S1.

### 2.2 Collins Test

The Collins Test assesses paw preference by quantifying forelimb use in a forced single-pellet retrieval task.^40^ Rats (20 females, 10 males) were placed in a custom chamber (11.4 cm height × 6.4 cm depth × 3.8 cm width) with a narrow frontal slot permitting the use of only one forepaw (Fig. 1A). Prior to testing, rats were habituated to sugar pellets (*Dustless Precision Pellets from Bio-Serv, Flemington, NJ, USA)* in their home cages and to the apparatus for three consecutive days; paw use was not recorded during habituation. A 24-hour mild food restriction (15 g/day) was imposed before testing. On test day, each rat was given 11 retrieval opportunities. The forepaw used for each retrieval was recorded, and paw preference was determined as the percentage of right vs. left paw use. Sessions were valid if ≥10 trials were completed; animals not meeting this threshold were retested the following day. This trial number was chosen based on prior literature, which shows reliable laterality estimates from 8–12 trials.^48, 49^ Extended sessions (>50 trials) were avoided to reduce fatigue and stress. Exploratory analyses confirmed that laterality indices stabilized by ∼10 trials, supporting the adequacy of 11. Chamber dimensions were not adjusted by age, as both adolescent and adult rats completed the task without difficulty.

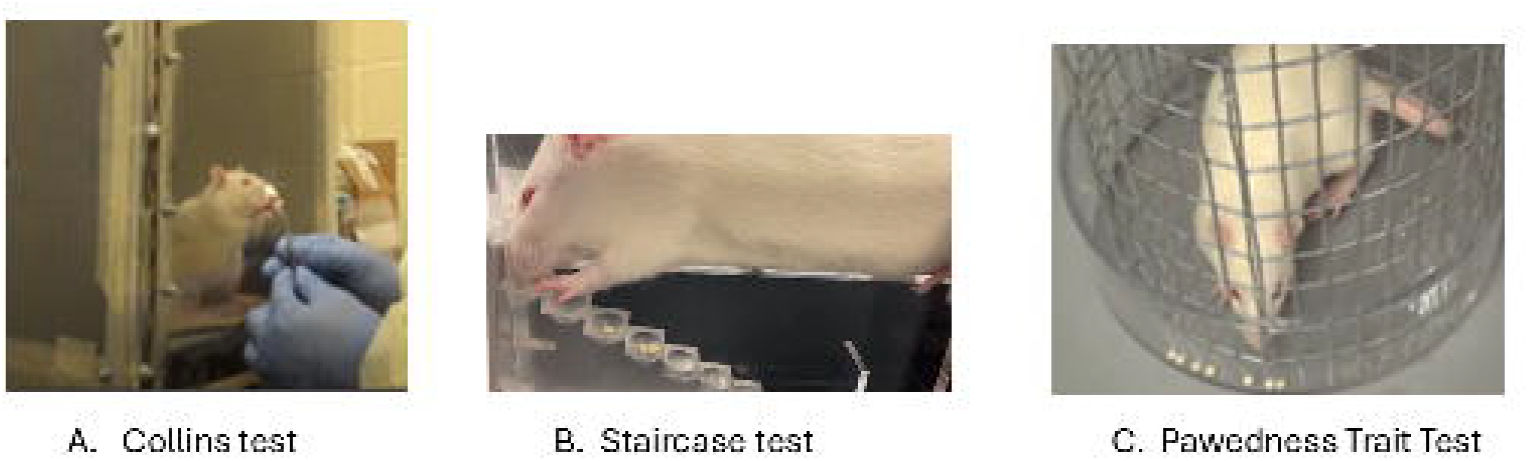

### 2.3 Staircase Test

The Staircase Test evaluates paw preference and skilled motor coordination through pellet retrieval from bilateral stepped platforms.^41^ The apparatus (Model 80300A, Lafayette Instrument, IN, USA; 36 × 12 cm) contained two staircases with sugar pellets placed on seven progressively higher steps (Fig. 1B). Prior to testing, rats underwent two habituation sessions (5 min and 10 min, with pellet exposure) followed by a short training phase with pellets placed at progressively higher steps. Testing comprised five 5-min sessions, with the final two including unilateral pellet placement to assess forced use. Performance during habituation/training was excluded from analysis. Paw preference was quantified by the number of pellets retrieved with each paw and the lowest step successfully reached. A dataset was considered valid if ≥10 retrievals per side were achieved across the five sessions. Rats were food-restricted 24 hours before testing (≥90% body weight maintained). Apparatus dimensions were constant across age groups; younger rats (6 weeks) successfully accessed all steps, confirming body size did not confound performance.

### 2.4 Pawedness Trait Test (PaTRaT)

The PaTRaT evaluates spontaneous paw preference in a free-choice context.^42^ Rats were confined in a 40 cm-long transparent mesh cylinder (17.5 cm diameter; 1.1 × 2.2 cm mesh openings) with an external reward holder (4 × 2 cm) at the base, positioned on a stable platform (50 × 75 cm) (Fig. 1C). The apparatus was obtained from Maze Engineer (Chicago, IL, USA). Pretraining included two 10-min familiarization sessions and two daily 15-min motivation sessions (spaced 4 h apart) with pellets placed near the mesh. Laterality data were collected only during the formal 10-min test session. Ten sugar pellets were distributed in the external reservoir, and each retrieval was scored as left, right, or both paws. Sessions were valid with ≥8 retrievals; if this criterion was not met, a repeat session was conducted after renewed habituation. Standard apparatus dimensions were used for all age groups. Observations confirmed that both younger and older rats could perform the task without restriction. Following testing, animals were returned to their home cages with unrestricted food and water.

### 2.5 Statistical Analysis

All analyses were conducted in GraphPad Prism v10 (GraphPad Software, CA, USA) and IBM SPSS v29 (IBM Corp., Armonk, NY, USA). Behavioral scoring was performed by a trained experimenter blinded to age and sex. To confirm reliability, 20% of sessions were independently re-scored by a second blinded rater; inter-rater agreement exceeded 95%.

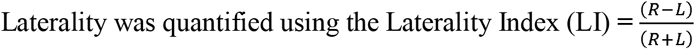

where *R* and *L* represent the number of successful right - and left-paw retrievals, respectively. LI values range from –1 (exclusive left paw use) to +1 (exclusive right paw use), with 0 indicating no preference. For consistency with literature, the term *LI* is used throughout the article.

Laterality was quantified using the Laterality Index (LI), which is mathematically equivalent to the Coefficient of Asymmetry (CA) when expressed as a proportion, as noted in some previous literatures.^13,18^

#### Classification Criteria

To classify paw preference, we adopted thresholds from prior rodent laterality studies. For the Collins and PaTRaT tests, LI –1≤ –0.4 indicated left-pawed, LI +1 ≥ +0.4 indicated right-pawed, and –0.4 < LI < +0.4 indicated ambidextrous.^50, 51^ A cutoff of ±0.4 has been widely used because it represents a moderate degree of asymmetry that exceeds normal within-subject variability while avoiding over-classification of weak preferences. For the Staircase Test, a narrower threshold of ±0.15 was applied, consistent with its design, where the bilateral pellet distribution yields more sensitive indices of side bias.^41^ PaTRaT raw scores (–4 to +4) were normalized where LI –1≤ –0.4 indicated left-pawed, LI +1 ≥ +0.4 indicated right-pawed, and –0.4 < LI < +0.4 indicated ambidextrous LI scale to harmonize classification across paradigms.^52^

#### Agreement Analysis

Inter-test reliability was quantified using Fleiss’ κ across all three tests, and pairwise Cohen’s κ with 95% confidence intervals. Categorical data were derived from LI thresholds, and bootstrap resampling (1,000 iterations) was used to compute confidence intervals. Interpretation followed Landis and Koch (1977): <0 = poor; 0.01–0.20 = slight; 0.21–0.40 = fair; 0.41–0.60 = moderate; 0.61–0.80 = substantial; 0.81–1.00 = almost perfect agreement.^53^

#### Analysis of Variance

For animals tested across all paradigms (*n* = 30), repeated-measures ANOVA was used to compare LI values between the Collins, Staircase, and PaTRaT tests. Population-level differences in laterality strength were analyzed using repeated measures ANOVA comparing the Laterality Index (LI) with Sex (male, female) and Age (adolescent: 6 weeks; adult: ≥12 weeks) as between-subject factors. Effect sizes were reported as partial η^2^, and significance was set at *p* < 0.05.

#### Exploratory Analyses

To test the robustness of our classification approach, threshold cutoffs were varied in increments of 0.05. While the absolute number of animals assigned to each category shifted slightly, inter-test agreement (κ values) remained stable, confirming that conclusions were not dependent on any single cutoff. Thresholds were also compared against LI histograms (Fig. 2A–C), which showed natural inflection points near the chosen values, further supporting their validity.

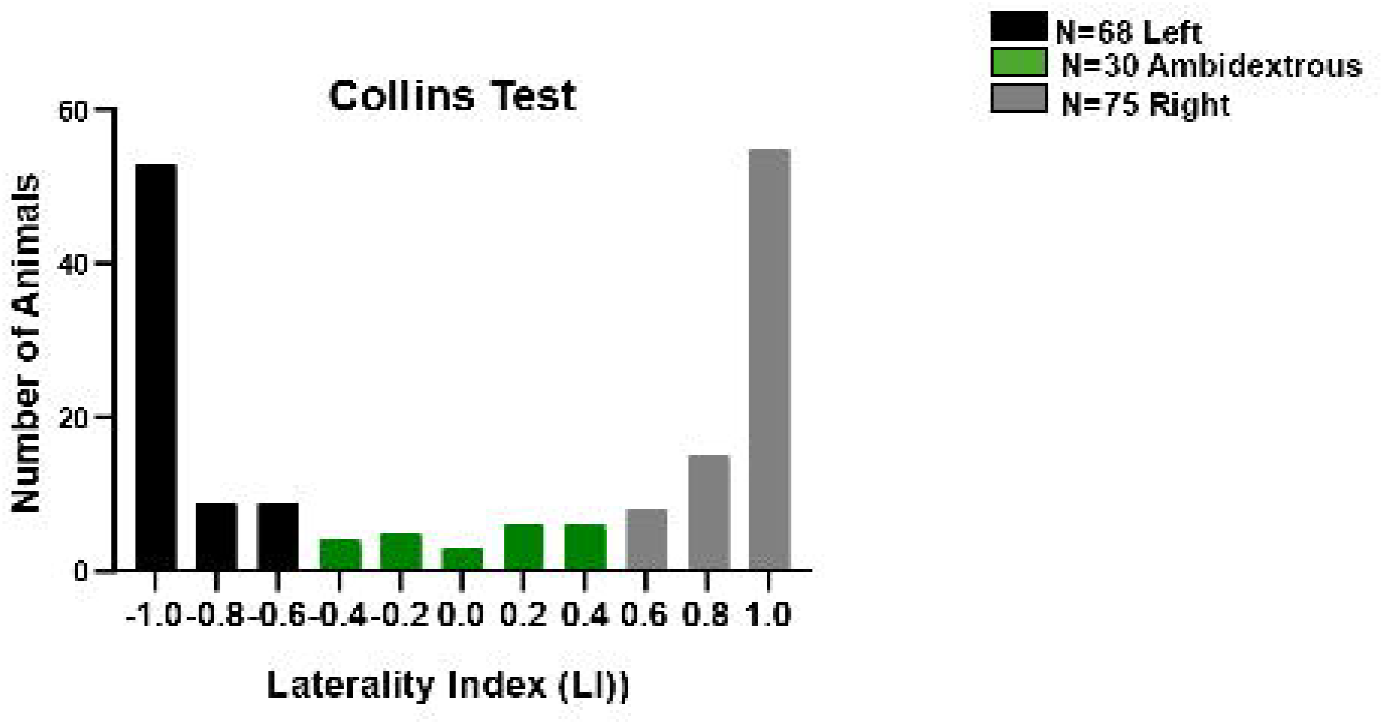

We acknowledge that categorical thresholds, while useful for between-study comparisons, are not universally standardized. Our choice of ±0.4 for Collins and PaTRaT reflects a conservative criterion that balances sensitivity and specificity, whereas ±0.15 for Staircase Test reflects its greater trial count and sensitivity to subtle asymmetries. Future studies may benefit from analyzing LI as a continuous measure rather than imposing arbitrary cutoffs, thereby improving reproducibility and allowing finer-grained analyses of individual variability.

## 3. Results

### 3.1 Distribution of Paw Preference in a Larger Cohort Assessed with the Collins Test

To examine population-level patterns of laterality, the Collins Test was applied to a larger cohort (*n* = 173). LI distributions revealed that most rats exhibited a lateralized preference, with relatively few classified as ambidextrous (Fig. 2). Of these, 75 were right-pawed, 68 left-pawed, and 30 ambidextrous. This corresponds to 43% right-pawed, 39% left-pawed, and 17% ambidextrous. These proportions are consistent with prior reports. For example, Manns et al. (2021) reported ∼45% right-pawed, ∼35% left-pawed, and ∼20% ambidextrous in a meta-analysis of rodents.^9^ Our cohort closely mirrors this distribution, reinforcing the robustness of the LI classification scheme and confirming that paw preference distributions generalize reliably from smaller to larger populations

### 3.2 Consistency of Paw Preference Across Behavioral Tests

Paw preference was assessed in the same cohort of rats (*n* = 30) using the Collins Test (Fig. 3A), Staircase Test (Fig. 3B), and PaTRaT (Fig. 3C). LI values were normalized across tests to enable direct comparison. Repeated measures ANOVA revealed no significant differences in LI among the three assays (*p* = 0.801), indicating that the overall strength of lateralization was consistent across tasks (Fig. 3D). Fleiss’ κ confirmed substantial inter-test agreement (κ = 0.761, *p* = 3.93 × 10□^12^), demonstrating that the three paradigms yield convergent results at the group level.

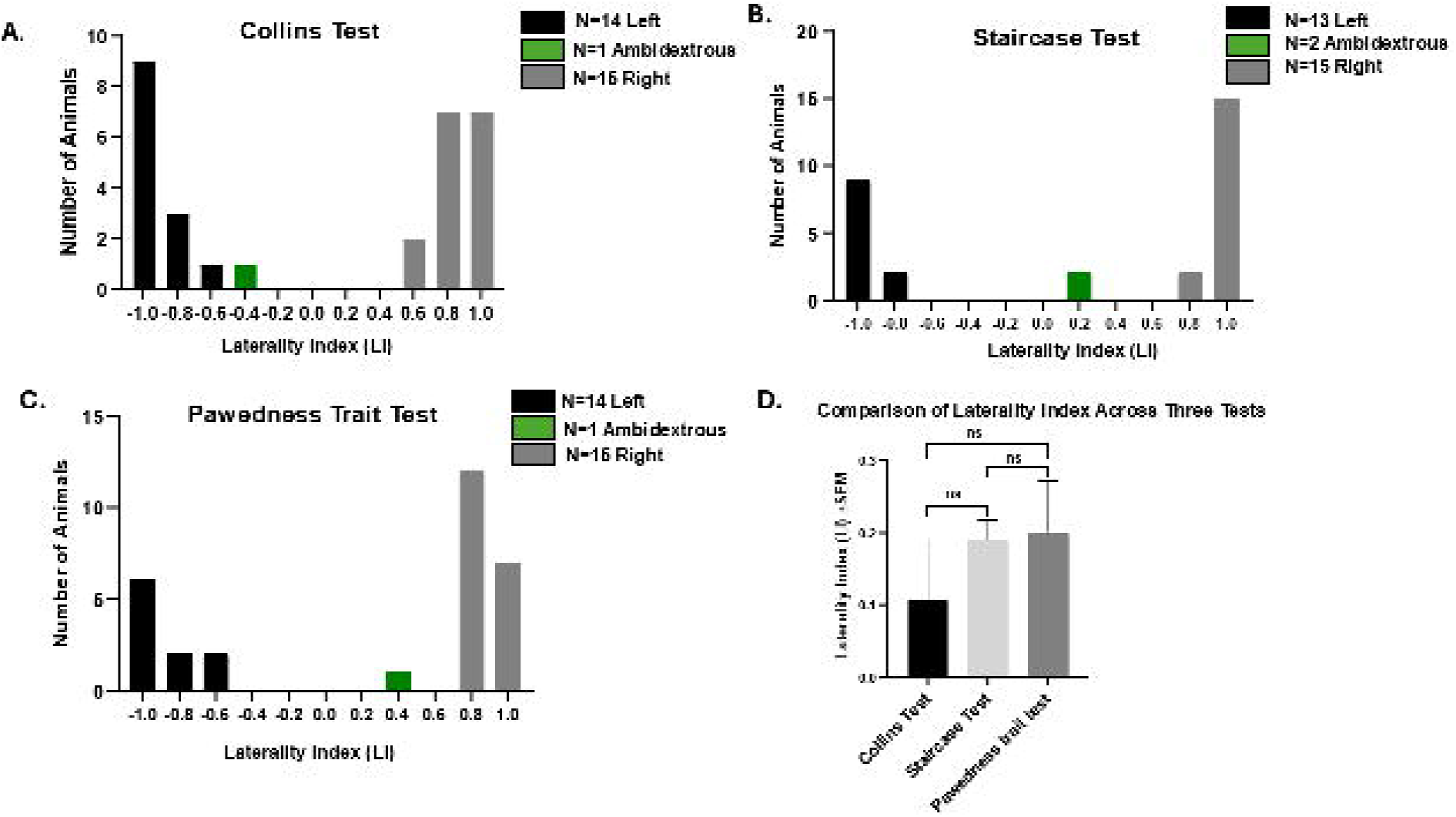

Pairwise agreement analyses are presented in Table 2. Agreement between the Collins and Staircase Tests was substantial for Cohen’s kappa (κ = 0.75, 95% CI [0.54–0.94]), indicating close concordance. Agreement between the Collins and PaTRaT was weak (κ = 0.18, 95% CI [– 0.18–0.52]), while Staircase vs. PaTRaT agreement was fair (κ = 0.27, 95% CI [–0.05–0.58]).

**Table 1.**
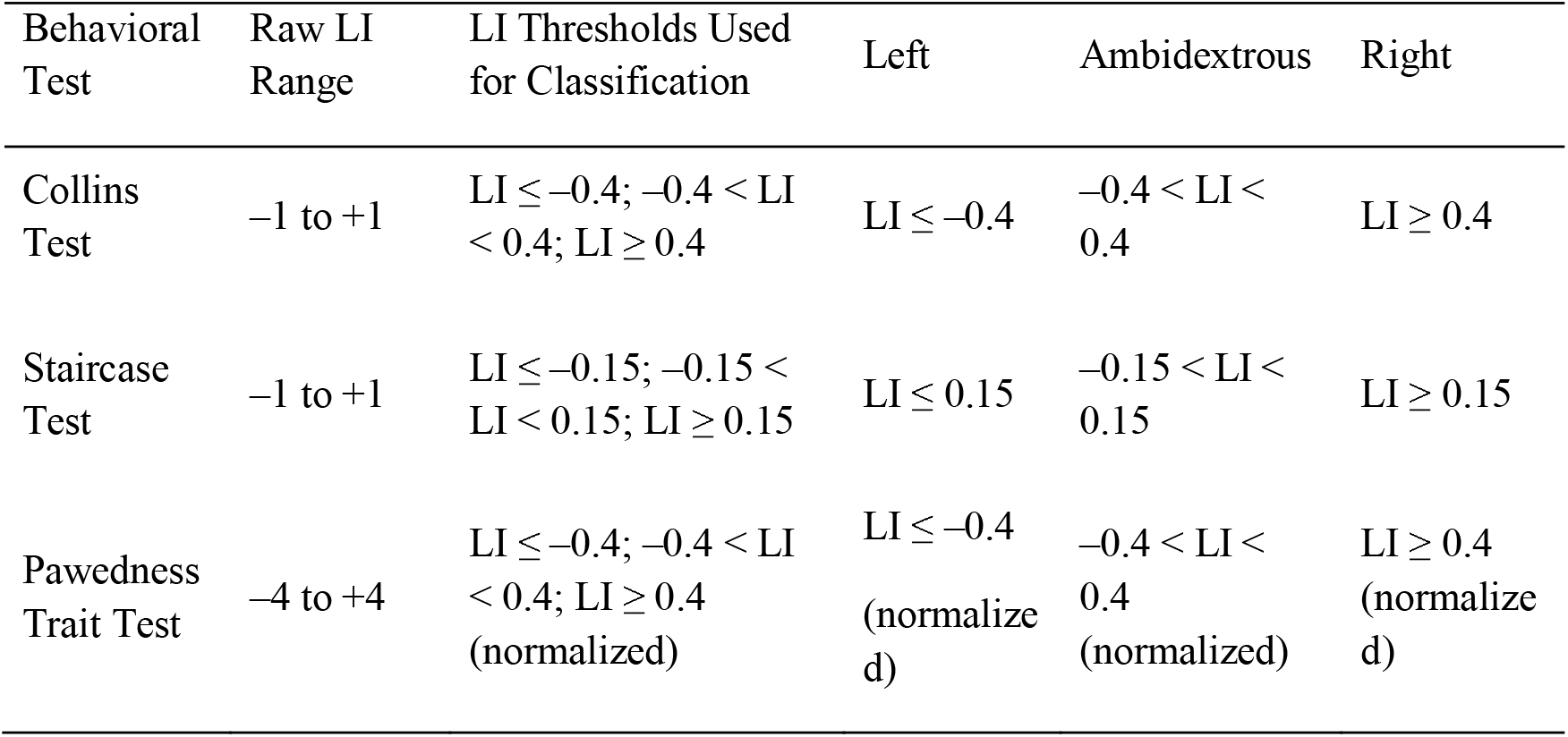
Classification criteria for paw preference across behavioral tests.

| Behavioral Test | Raw LI Range | LI Thresholds Used for Classification | Left | Ambidextrous | Right |
| --- | --- | --- | --- | --- | --- |
| Collins Test | $-1$ to $+1$ | $LI \leq -0.4$; $-0.4 < LI < 0.4$; $LI \geq 0.4$ | $LI \leq -0.4$ | $-0.4 < LI < 0.4$ | $LI \geq 0.4$ |
| Staircase Test | $-1$ to $+1$ | $LI \leq -0.15$; $-0.15 < LI < 0.15$; $LI \geq 0.15$ | $LI \leq 0.15$ | $-0.15 < LI < 0.15$ | $LI \geq 0.15$ |
| Pawedness Trait Test | $-4$ to $+4$ | $LI \leq -0.4$; $-0.4 < LI < 0.4$; $LI \geq 0.4$ (normalized) | $LI \leq -0.4$ (normalized) | $-0.4 < LI < 0.4$ (normalized) | $LI \geq 0.4$ (normalized) |

**Table 2.** Pairwise agreement across paw preference tests.

| Test Pair | Cohen's ($\kappa$) | 95% CI | N |
| --- | --- | --- | --- |
| Collins Test vs Staircase Test | 0.747 | 0.535 to 0.935 | 30 |
| Collins Test vs Pawedness Trait Test | 0.182 | -0.175 to 0.521 | 30 |
| Staircase Test vs Pawedness Trait Test | 0.267 | -0.054 to 0.576 | 30 |

These results suggest that although Collins and Staircase yield comparable classifications, PaTRaT introduces more variability and may capture distinct aspects of spontaneous paw use *(see Supplementary Table S2 for full analysis)*.

### 3.3 Stability of Paw Preference Across Age and Sex

Subgroup analyses were conducted using data from the Collins Test (Fig. 4). Age and sex did not significantly influence paw preference outcomes. Fig. 4, bar graph comparing left and right paw use in younger (6–9 weeks; n = 90) and older (12–36 weeks; n = 83) rats. Two-way ANOVA (Sex × Age) revealed no significant main effect of age [F (1, 378) = 0.146, p = 0.702] and no significant interaction between age and sex [F (3, 378) = 1.228, p = 0.299], indicating that laterality index (LI) remains stable across developmental stages and sexes. Bars represent mean ± SEM. Comparisons were performed using two-way ANOVA; “ns” denotes non-significant differences (p ≥ 0.05) *(see Supplementary Table S3 and S4)*.

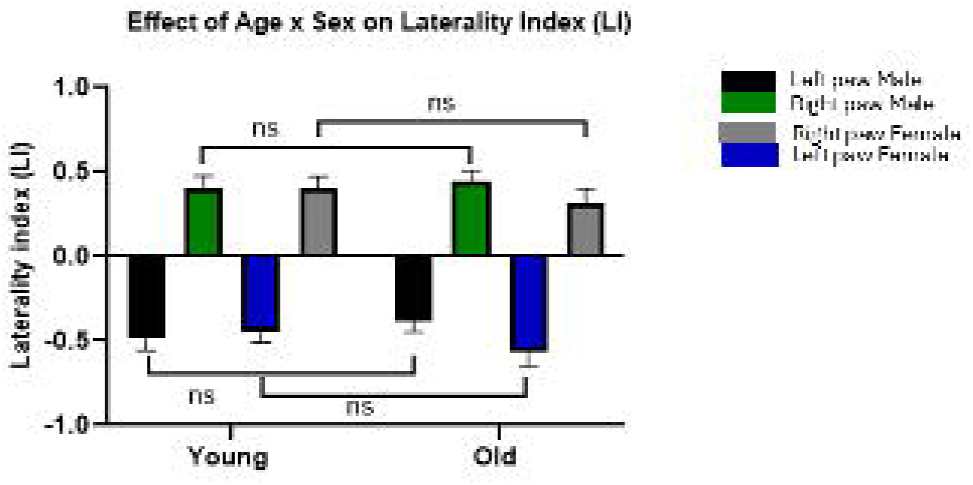

## 4. Discussion

This study demonstrates that three widely used behavioral assays, the Collins, Staircase, and PaTRaT tests yield convergent measures of lateralized forelimb use in Sprague Dawley rats. Despite differences in task design, ranging from forced single-pellet retrieval to skilled reaching and spontaneous free-choice behavior, all three assays produced consistent laterality indices (LI). Repeated measures ANOVA confirmed no significant differences in LI across tasks, and Fleiss’ κ analysis indicated substantial overall agreement. These findings support the conclusion that paw preference represents a robust and trait-like behavioral phenotype rather than a transient state influenced by context or motivation. Importantly, our work provides evidence for the validity of standardized LI thresholding across paradigms, enhancing reproducibility and comparability in preclinical research.^9^

### 4.1 Neural Basis and Translational Relevance

The behavioral consistency observed here likely reflects underlying asymmetries in sensorimotor circuits, particularly within motor cortex and basal ganglia pathways, which show lateralized organization in both rodents and humans.^21, 28, 54-56^ Dopaminergic signaling is also frequently asymmetric and plays a central role in motor laterality, with relevance to conditions such as stroke, cerebral palsy, traumatic brain injury, and lateralized cognitive functions including language.^32, 43, 57-60^ Although this study focused on behavioral endpoints, the validated assays provide accessible readouts of hemispheric dominance that can be integrated with neuroimaging, electrophysiological, or molecular approaches. Such multimodal studies could help map the anatomical and neurochemical substrates of lateralization.

### 4.2 Influence of Biological Factors

No significant differences in paw preference were detected across age or sex, supporting the view that paw preference is a stable trait in healthy rats. However, the potential plasticity of lateralization under specific conditions remains an open question. Prior works have shown that environmental enrichment, brain injury, and pharmacological manipulation can alter lateralized behaviors.^61-65^ Future research should therefore examine whether paw preference can be modified during recovery or rehabilitation, particularly in the context of therapies aimed at promoting neuroplasticity and neurogenesis.

### 4.3 Methodological Considerations

While inter-test agreement was generally strong, PaTRaT showed weaker concordance with the Collins and Staircase Tests. This variability may reflect the spontaneous, less structured nature of PaTRaT, which is prone to observer bias but also captures habitual paw use in a naturalistic context. In contrast, Collins and Staircase offer clearer endpoints but may be more influenced by food restrictions, task fatigue, and training effects. Methodological refinements such as automated video-based pose estimation (e.g., DeepLabCut) could reduce observer bias, increase throughput, and allow for more fine-grained kinematic analysis across all three paradigms.^66, 67^ Additionally, longitudinal studies are warranted to assess intra-individual stability and environmental sensitivity, strengthening the utility of paw preference as a biomarker for motor asymmetry.

Although our data shows convergence across tests, the relatively weak agreement involving the PaTRaT warrants a more cautious interpretation of cross-paradigm consistency. This divergence may reflect differences in task structure, spontaneous versus directed behavior, or motivational variability, all of which can influence paw use.^17, 38^ Additionally, the study focuses exclusively on behavioral outcomes without incorporating neurobiological correlates, limiting mechanistic insight into the neural substrates of lateralization.^19, 21^

### 4.4 Comparative Utility of the Three Tests

Table 3 summarizes the strengths, limitations, and potential applications of each assay. The Collins Test remains highly reliable, efficient, and historically validated; the Staircase Test is particularly sensitive to fine motor deficits and rehabilitation models; and PaTRaT captures spontaneous, low-stress behavior, making it suitable for screening or correlating motor laterality with emotional and cognitive traits. Together, these assays provide complementary insights and can be strategically selected based on the scientific question, desired sensitivity, and resource constraints.

**Table 3.**
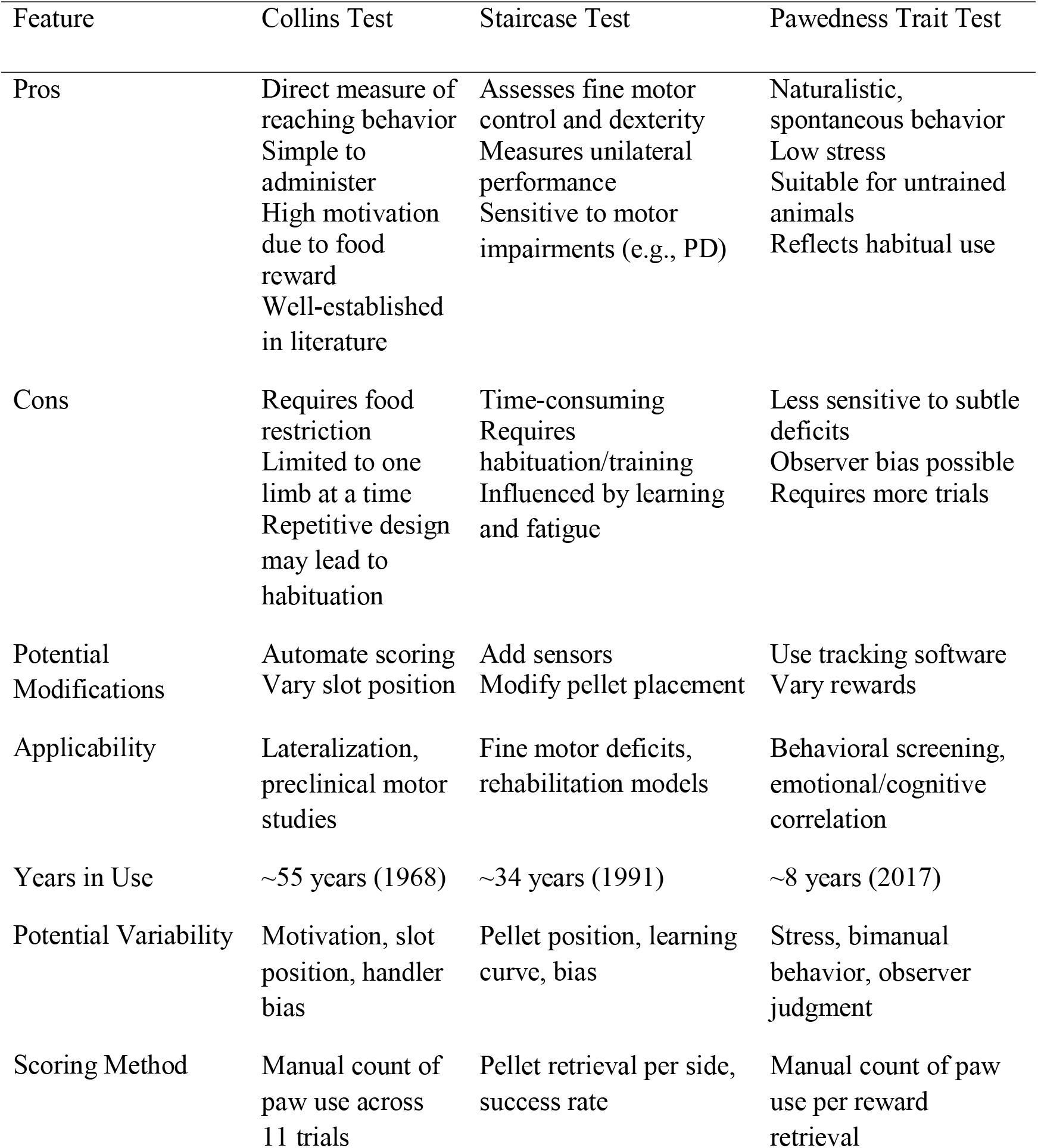
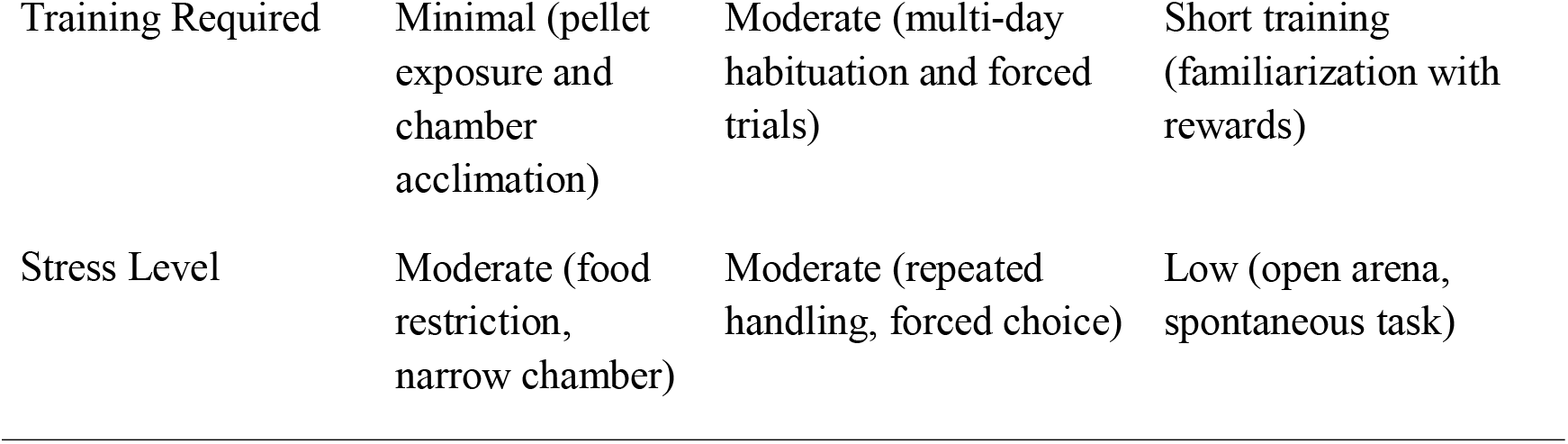
Comparative analysis of three behavioral tests for assessing paw preference in rats.

| Feature | Collins Test | Staircase Test | Pawedness Trait Test |
| --- | --- | --- | --- |
| Pros | Direct measure of reaching behavior<br>Simple to administer<br>High motivation due to food reward<br>Well-established in literature | Assesses fine motor control and dexterity<br>Measures unilateral performance<br>Sensitive to motor impairments (e.g., PD) | Naturalistic, spontaneous behavior<br>Low stress<br>Suitable for untrained animals<br>Reflects habitual use |
| Cons | Requires food restriction<br>Limited to one limb at a time<br>Repetitive design may lead to habituation | Time-consuming<br>Requires habituation/training<br>Influenced by learning and fatigue | Less sensitive to subtle deficits<br>Observer bias possible<br>Requires more trials |
| Potential Modifications | Automate scoring<br>Vary slot position | Add sensors<br>Modify pellet placement | Use tracking software<br>Vary rewards |
| Applicability | Lateralization, preclinical motor studies | Fine motor deficits, rehabilitation models | Behavioral screening, emotional/cognitive correlation |
| Years in Use | ~55 years (1968) | ~34 years (1991) | ~8 years (2017) |
| Potential Variability | Motivation, slot position, handler bias | Pellet position, learning curve, bias | Stress, bimanual behavior, observer judgment |
| Scoring Method | Manual count of paw use across 11 trials | Pellet retrieval per side, success rate | Manual count of paw use per reward retrieval |
| Training Required | Minimal (pellet exposure and chamber acclimation) | Moderate (multi-day habituation and forced trials) | Short training (familiarization with rewards) |
| Stress Level | Moderate (food restriction, narrow chamber) | Moderate (repeated handling, forced choice) | Low (open arena, spontaneous task) |

### 4.5 Limitations

I. Sample size and design: Although the inter-test cohort (n = 30) enabled within-subject comparisons across the three behavioral paradigms, this modest sample size may limit statistical power for detecting subtle differences, especially in age- and sex-based subgroups. Nonetheless, it exceeds the sample sizes used in most prior validation studies (typically n = 12–20 per group) and was supplemented by a larger Collins-only cohort (n = 173) to confirm population-level distributions.^42, 45, 68-70^ Future studies employing fully counterbalanced designs with larger cohorts, randomized task orders, and extended washout periods between tests will help mitigate potential carry-over or learning effects.
II. Trial numbers and task duration: To balance reliability with animal welfare, the Collins Test was limited to 11 retrieval attempts per animal. This reduced handling time and prevented fatigue or stress associated with longer sessions. Exploratory analyses indicated that laterality indices stabilized by approximately 10 trials; however, this may not capture finer gradations of preference observed in longer protocols (≥ 50 trials).^40, 48, 49^ A systematic evaluation of how trial number influences classification stability would clarify minimum trial requirements for reliable laterality estimation.
III. Threshold-based classification: The use of fixed Laterality Index cutoffs (± 0.4 for Collins/PaTRaT and ± 0.15 for Staircase) allowed comparability with existing literature but introduced categorical rigidity. Although sensitivity analyses demonstrated that inter-test agreement remained stable across nearby cutoffs, dividing continuous behavior inevitably obscures subtle inter-individual differences. Moreover, normalization of PaTRaT raw scores (–4 to +4) to the –1 to +1 LI scale added a transformation step that may slightly distort distributional properties. The current dataset was analyzed primarily at the subject-level LI, which is typical for this literature.^17, 71, 72^ Future studies could be designed to use mixed-model or hierarchical analyses to capture both direction and magnitude of lateralization within individuals.^73, 74^
IV. Manual scoring and observer variability: Another limitation is the reliance on manual scoring. Although all raters were trained to criterion and followed standardized protocols, inter-rater reliability was not formally quantified in this study. Although 20 % of sessions were re-scored for reliability (> 95 % concordance), formal inter-rater statistics were not computed for the full dataset. Integration of automated pose-estimation platforms such as DeepLabCut or SimBA could enhance reproducibility, reduce observer bias, and allow high-resolution kinematic analyses of reach trajectories and digit use.^70^
V. Strain and Environmental generalizability: All experiments were conducted on Sprague Dawley rats, which are commonly used but may differ from other strains in cortical organization, stress reactivity, and motivational profiles. Results may therefore not generalize to Long-Evans, Wistar, or Lister Hooded rats without further validation.^38, 39, 75-77^ Additionally, environmental factors such as lighting, cage enrichment, and food restriction regimes were carefully standardized but may influence task motivation and lateralization outcomes in other laboratories. Cross-laboratory replication and multi-strain validation will be essential for establishing universal reference ranges for paw preference metrics.
VI. Scope and correlational constraints: The present work focused exclusively on behavioral outcomes. Although these tests are presumed to reflect hemispheric motor dominance, direct neuroanatomical or neurochemical correlations (e.g., asymmetric striatal Tyrosine hydroxylase (TH), Vesicular monoamines transferase 2 (VMAT2), or dopa decarboxylase (DDC) expression) were not assessed.^19, 20, 79^ Future studies integrating histological, electrophysiological, or imaging endpoints would provide critical insight into the neural substrates underlying behavioral lateralization and its modulation by disease or intervention.

## 5. Conclusion

This study demonstrates that the Collins, Staircase, and Pawedness Trait Tests yield convergence and reliable measures of paw preference in Sprague Dawley rats. Substantial inter-test agreement confirms that paw preference is a stable, trait-like behavioral feature unaffected by age or sex.

By standardizing LI thresholds and experimental procedures, this work resolves prior inconsistencies and establishes a reproducible framework for assessing motor lateralization. These validated assays complement one another with the Collins and Staircase to capture task-directed precision, while PaTRaT reflects spontaneous behavior allowing flexible applications across diverse experimental designs. Together, they provide validated behavioral tools for modeling hemispheric specialization, investigating asymmetric dopaminergic function, and probing selective vulnerability in neurological and neurodegenerative disorders. Future studies integrating neurochemical or imaging correlates will further clarify the neural mechanisms linking behavioral laterality with hemispheric motor dominance.

## Abbreviations

LI: Laterality Index
PaTRaT: Pawedness Trait Test
PD: Parkinson’s disease

## Ethics statement

We confirm that we have read the Journal’s Position on Ethics in Publishing and that this manuscript is consistent with those guidelines.

## CRediT authorship contribution statement

D. Pokharel: Conceptualization, methodology, validation, writing and editing, data collection and analysis. K. Le: Contributed to protocol development and execution of the study. D. Beligala: Provided the foundational concepts, reviewing and editing. T. Subramanian: Conceptualization, resources, editing, supervision, project administration, and funding acquisition. K. Venkiteswaran: Additional contributions in resources, supervision, and project administration.

## Acknowledgements

We would like to acknowledge Dr. Sadik Khuder, PhD, Professor of Medicine, Statistics, and Public Health at The University of Toledo, Toledo, OH 43614; Dr. Daniel Leventhal, MD, PhD, Professor of Neurology at the University of Michigan, Ann Arbor, MI 48109; Dr. James Burkett, PhD, Professor of Neurosciences and Psychiatry at the University of Toledo College of Medicine and Life Sciences, Toledo, OH 43614; and Dr. Tim Gilmour, PhD, Professor at John Brown University, Siloam Springs, AR 72761, for their valuable feedback and guidance.

## Declaration of generative AI and AI-assisted technologies

During the preparation of this work, the authors used Perplexity AI to assist with grammar and language editing. Following its use, the authors reviewed and revised the content as necessary and took full responsibility for the integrity and accuracy of the final manuscript.

## Data availability

Data will be available upon request.

## Appendix

**Supplemental Table S1.** Summary of Rat Cohorts and Tests Administered:

| Cohort | n | Age Range | Sex Distribution | Tests Administered |
| --- | --- | --- | --- | --- |
| Core Inter-Test Group | 30 | 12–48 wks | 20 females, 10 males | Collins, Staircase, PaTRaT |
| Young Age Comparison | 90 | 6–9 wks | Mixed | Collins Test only |
| Older Age Comparison | 83 | 12–36 wks | Mixed | Collins Test only |
| Male Sex Comparison | 93 | 6–48 wks | All male | Collins Test only |
| Female Sex Comparison | 80 | 6–48 wks | All female | Collins Test only |

